# Tumor regression mediated by oncogene withdrawal or erlotinib stimulates infiltration of inflammatory immune cells in EGFR mutant lung tumors

**DOI:** 10.1101/254847

**Authors:** Deborah Ayeni, Braden Miller, Alexandra Kuhlman, Ping-Chih Ho, Camila Robles-Oteiza, Mmaserame Gaefele, Stellar Levy, Fernando J. de Miguel, Curtis Perry, Tianxia Guan, Daniel Zelterman, Robert Homer, Zongzhi Liu, Susan Kaech, Katerina Politi

**Author notes:** To whom correspondence should be addressed: Katerina Politi, PhD, Department of Pathology, Yale Cancer Center, Yale University School of Medicine, 333 Cedar Street, SHM-I 234D, New Haven, CT 06510 USA. The authors have the following conflicts to disclose: Research funding from AstraZeneca (KP, SMK), Roche (KP, SMK), Kolltan (KP). Consulting/Advisory Role honoraria from AstraZeneca (KP), Merck (KP), Novartis (KP), Tocagen (KP) Royalties in IP licensed from MSKCC to Molecular MD (KP).

## Abstract

Epidermal Growth Factor Receptor (EGFR) tyrosine kinase inhibitors (TKIs) like erlotinib are effective for treating patients with EGFR mutant lung cancer; however, drug resistance inevitably emerges. Approaches to combine immunotherapies and targeted therapies to overcome or delay drug resistance have been hindered by limited knowledge of the effect of erlotinib on tumor-infiltrating immune cells. Using mouse models, we studied the immunological profile of mutant *EGFR*-driven lung tumors before and after erlotinib treatment. We found that erlotinib triggered the recruitment of inflammatory T cells into the lungs. Interestingly, this phenotype could be recapitulated by tumor regression mediated by deprivation of the EGFR oncogene indicating that tumor regression alone was sufficient for these immunostimulatory effects. Erlotinib treatment also led to increased maturation of myeloid cells and an increase in CD40+ dendritic cells. Our findings lay the foundation for understanding the effects of TKIs on the tumor microenvironment and highlights potential avenues for investigation of targeted and immuno-therapy combination strategies to treat EGFR mutant lung cancer.

## Introduction

*EGFR* mutations are found in 10-15% of lung adenocarcinomas in the US and are enriched in tumors from never or former smokers [1]. Lung adenocarcinoma-associated mutations in exons encoding the tyrosine kinase domain of this receptor most commonly include either deletion of a four amino acid motif (LREA) in Exon 19 of *EGFR* or a point mutation in Exon 21, which substitutes Arginine for Leucine at position 858 (L858R) [2]. These mutations confer sensitivity to EGFR tyrosine kinase inhibitors (TKIs) such as erlotinib, gefitinib and afatinib, current standard of care therapies for the treatment of this subset of lung cancer. However, drug resistance inevitably develops on average after 12 months of treatment [3, 4]. In more than 50 percent of cases, acquired resistance to erlotinib is driven by a second site mutation in *EGFR*, T790M [3, 5], which alters the affinity of the receptor for ATP and as a consequence to the drugs [6]. Ongoing promising efforts to overcome and/or delay the onset of resistance include novel 3^rd^ generation TKIs that specifically inhibit mutant EGFR (and spare wild-type EGFR) [7] and dual EGFR blockade using an anti-EGFR antibody/TKI combination [8]. Even with these improvements, however, none of the therapies are curative [9]. Therefore, demands for novel therapeutic approaches are high.

Recent advances show that targeting the immune system is a useful approach to treating lung cancer. Mounting evidence suggests that tumors stimulate the establishment of an immunosuppressive microenvironment to evade the immune system by facilitating tumor-infiltrating T cells to display an exhausted phenotype, [10] such that they are unable to proliferate and produce pro-inflammatory cytokines [11, 12]. Agents that target inhibitory molecules (e.g. PD-1, CTLA4) on T cells and/or their cognate ligands (e.g. PD-L1) on tumor and immune infiltrating cells have shown promising results in treating lung cancers in clinical trials [13]. In a subset of patients with advanced, pretreated lung cancer, treatment with nivolumab, an anti-PD-1 blocking antibody, objective responses were observed in approximately 20% of patients, which in some cases was prolonged [14]. Furthermore, patients with previously treated, PD-L1 positive lung tumors receiving 10mg/kg pembrolizumab (another PD-1 targeting antibody) had significantly longer overall survival compared to patients treated with docetaxel (17.3 months vs 8.2 months) [15]. Notably, patients with untreated, advanced NSCLC with at least 50% PD-L1 tumor expression treated with pembrolizumab had a significantly longer progression-free survival compared to platinum-based chemotherapies [16], demonstrating the efficacy of this agent in a first-line setting and supporting tumor expression of PD-L1 as a marker of response to immune checkpoint inhibitors. In the latter study patients with tumors harboring sensitizing mutations in *EGFR* were excluded. In fact, in a retrospective evaluation of patients treated with PD-1 or PD-L1 inhibitors, it was found that objective responses in patients with *EGFR*-mutant tumors was 3.6% compared to 23.3% in those with *EGFR* wild-type tumors [17]. This suggests that there is a lower response rate to the PD-1 axis inhibitors associated with this genetic alteration. In spite of this, there are clear indications that a subset of patients with *EGFR* mutant lung cancer benefit from these therapies [15, 18, 19]. Moreover, preclinical models demonstrate that the immune system plays an important role in modulating the growth of *EGFR* mutant tumors [20]. In one study evaluating the combination of erlotinib plus nivolumab, durable tumor regression in both treatment (TKI or chemotherapy) naïve and TKI-treated patients was reported [21] and there are several additional trials evaluating the efficacy of combining PD-1/PD-L1 inhibitors with EGFR TKIs [18]. However, toxicities have raised concerns that treating patients with EGFR TKIs with immune checkpoint inhibitors concurrently may not be the optimal approach to use these agents in combination. Given these findings, studies are necessary to understand the effects of EGFR TKIs on the tumor microenvironment and the immunological consequences of combining immune checkpoint inhibitors with EGFR TKIs.

Several studies have examined the effect of kinase inhibitors on the tumor immune microenvironment. The BRAF inhibitor vemurafenib, for instance, has been reported to increase intratumoral CD8^+^ T cell infiltrates [22], increase tumor associated antigens and improve effector function of cytotoxic T lymphocytes [23]. However, a subset of tumors resistant to vemurafenib exhibit features of T-cell exhaustion and reduced antigen presentation suggesting that these may be resistant to checkpoint inhibitors [24]. Similarly, in lung cancer cell lines, two studies have revealed that TKI treatment leads to down-regulation of tumor PD-L1 expression [25, 26]. Moreover, it has also been shown that erlotinib can impair T cell-mediated immune responses via suppression of signaling pathways downstream of EGFR critical for cell survival and proliferation [27]. Further supporting that erlotinib could have immunosuppressive effects on the immune system, erlotinib has been posited to down-regulate TNF-α mediated inflammation characteristic of psoriasis [28]. In addition, a recent study in mouse models of *EGFR* mutant lung cancer reported increased leukocyte infiltration and enhanced antigen-presenting capabilities after erlotinib treatment [29]. However, an investigation of whether these effects are mediated by erlotinib or tumor regression *in vivo* is lacking. To address this issue, we utilized a previously developed immunocompetent mouse model of *EGFR* mutant lung cancer [30] and tested the consequences of erlotinib or oncogene de-induction on the immune microenvironment.

## Results

### Increased inflammatory T cells following erlotinib treatment in EGFR mutant lung cancer mouse models

To evaluate the changes that occur in the immune microenvironment upon TKI treatment, *CCSP-rtTA; TetO-EGFR^L858R^* mice on a doxycycline diet were treated with erlotinib, an EGFR TKI, for a period of 2 weeks (**Figure 1A**). In tumor bearing mice after 2 weeks of erlotinib treatment, the disease is mostly undetectable by Magnetic Resonance Imaging (MRI) (**Supplementary Figure 1A**) and largely resolved histopathologically (**Supplementary Figure 1B**). At the end of the treatment, lung and spleen single cell suspensions were prepared and analyzed by flow cytometry. We compared the immune profiles of normal healthy lungs and lungs from tumor-bearing untreated and erlotinib-treated mice. To ensure that the effects observed were not due to the presence of doxycycline in the mouse diet, all of the mice, including controls were maintained on doxycycline for the same amount of time. We found a consistent reduction in the fraction of CD45^+^ immune cells and the absolute number of CD4^+^ and CD8^+^ T cells per gram of lung tissue in untreated tumor-bearing lungs that was reversed upon TKI treatment (**Figure 1B and Supplementary Figures 1C&D**).

**Figure 1.**
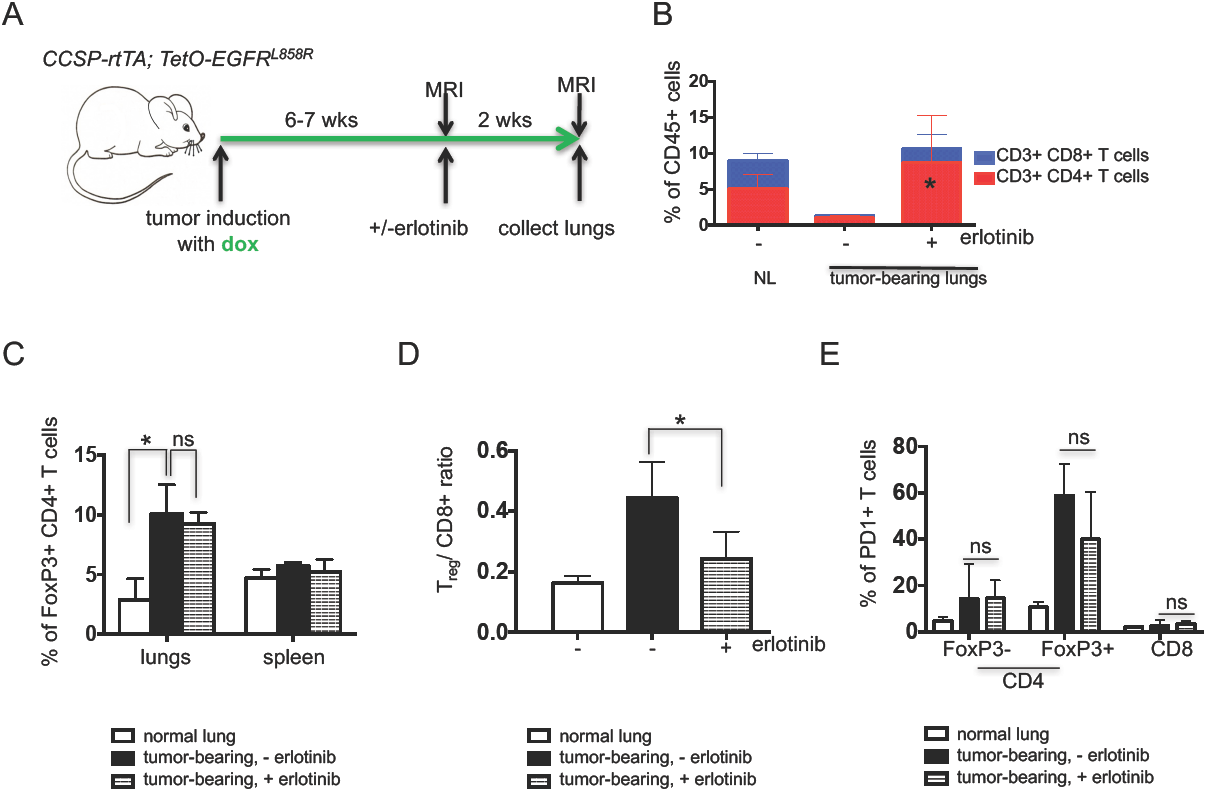
The immunosuppressive microenvironment in murine EGFR^L858R^ -induced lung adenocarcinomas is partially reversed by erlotinib. (**A**) Experimental outline of tumor induction and erlotinib treatment. *CCSP-rtTA; TetO-EGFR^L858R^* mice and littermate controls on a doxycycline diet (green arrow) for 6-7 weeks were treated with erlotinib or left untreated for 2 weeks. Infiltrating immune cells were analyzed by flow cytometry. Quantification of (**B**) CD4 and CD8 T cells (**C**) FoxP3 positive CD4 T cells (**D**) T_reg_/ CD8+ T cell ratio and (**E**) PD1 positive FoxP3- and FoxP3+ CD4 and CD8 T cells in the lungs (and spleens) of normal lung (NL) and tumor bearing *CCSP-rtTA; TetO-EGFR^L858R^* mice in the absence (-) and presence (+) of erlotinib for 2 weeks. Data are obtained from three independent experiments, (n=4-6 mice per group) *, P < 0.05 in student t test.

To determine whether there were any differences in the T-cells in tumor-bearing lungs indicative of an immunosuppressive microenvironment, we quantified regulatory T-cells present in the different conditions. We observed a significant increase in Foxp3^+^ regulatory T-cells (T_regs_) in the lungs of tumor-bearing mice regardless of erlotinib treatment (**Figure 1C and Supplementary Figure 1E**) suggesting that these immunosuppressive cells, which also may play a role in tissue repair, are retained even following erlotinib-mediated tumor regression. Despite the lack of a major shift in the proportion of T_regs_ in the erlotinib-treated lungs, the T_reg_/CD8^+^ T cell ratio decreased with erlotinib treatment, likely due to the increase in CD8^+^ T cells and indicative of a shift towards a more immunostimulatory microenvironment (**Figure 1D**). Interestingly, these T_regs_ retained a high level of PD-1 expression that was unchanged with erlotinib treatment (**Figure 1E and Supplementary Figure 1F**). To confirm these findings using an orthogonal approach, we used immunofluorescence to detect the tumor cell marker, cytokeratin, a pan T cell marker CD3, and the T_reg_ marker, Foxp3. We observed that erlotinib treatment induced infiltration of T cells into the lungs compared to untreated tumor-bearing lungs (**Supplementary Figure 1G**). Our quantification of Foxp3^+^ cells from these sections also revealed that there was no significant difference in their abundance between untreated and erlotinib-treated lungs (**Supplementary Figure 1H**). *In vitro* T-cell stimulation assays demonstrated that both CD4^+^ and CD8^+^ T cells showed increased production of the IFN-γ, TNF-α and IL-2 cytokines after erlotinib treatment indicative of an activated phenotype (**Figures 2A&B and Supplementary Figure 2A**). These results suggest the presence of an immunosuppressive microenvironment in the lungs of mice with *EGFR^L858R^* tumors, which is consistent with recent findings from a mouse model of *EGFR^Ex19del^* mutant lung cancer [20]. Erlotinib treatment leads to an increase in the numbers of lymphocytes, their higher cytokine production and a limited reduction in the proportion of T_regs_.

**Figure 2.**
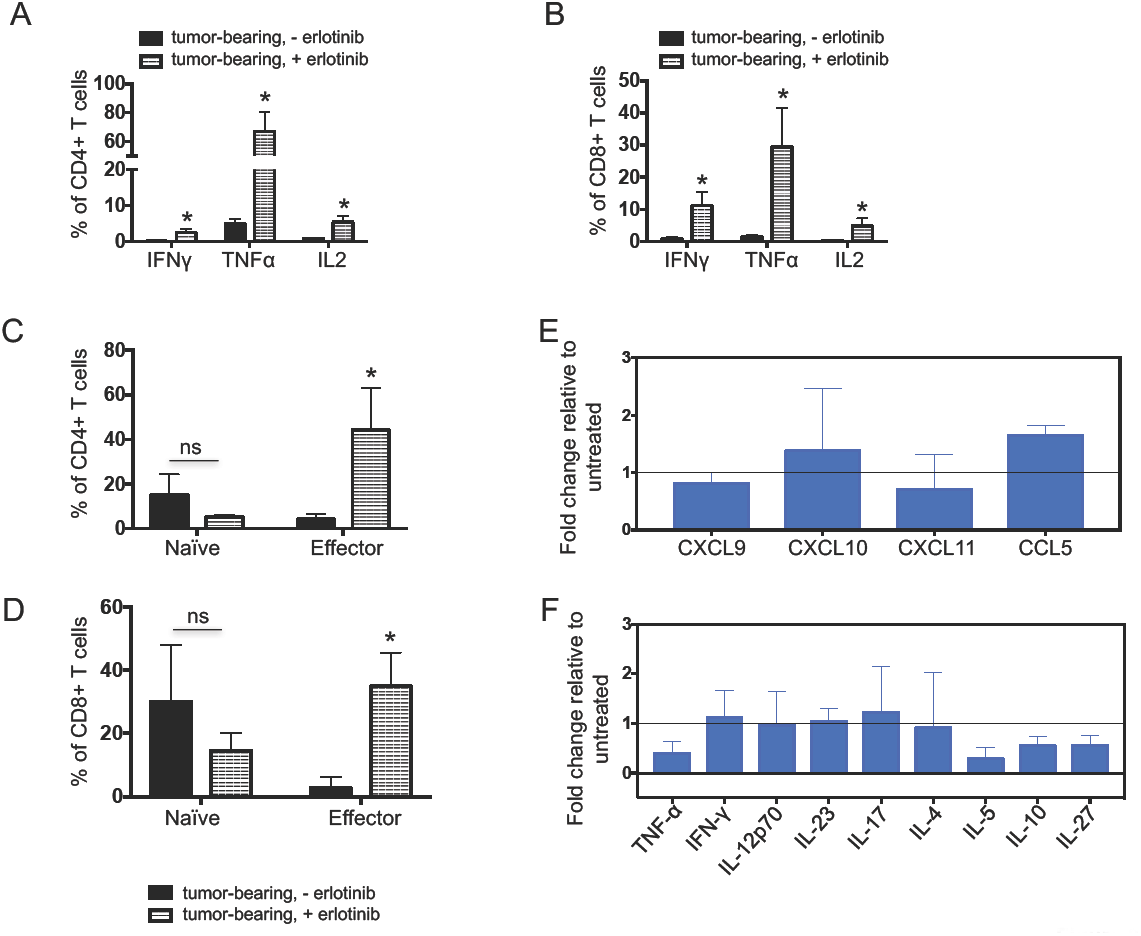
Increased production and presence of immunostimulatory cytokines following erlotinib treatment. Quantification of the levels of indicated effector cytokines from (**A**) CD4 T cells and (**B**) CD8 T cells after PMA/ionomycin stimulation and intracellular cytokine staining of cells in the lungs of tumor bearing *CCSP-rtTA; TetO-EGFR^L858R^* mice in the absence (-) and presence (+) of erlotinib for 2 weeks. Quantification of naïve and effector (**C**) CD4 and (**D**) CD8 T cells in lungs of *CCSP-rtTA; TetO-EGFR^L858R^* tumor bearing mice untreated or treated with erlotinib for 2 weeks. Graph representing different (**E**) chemokines and (**F**) cytokines screened in erlotinib treated whole lung tissue lysates normalized to levels in untreated lung lysates. Data are from three independent experiments, (n=4-6 mice per group) *, P < 0.05 in student t test.

To further study the properties of tumor-infiltrating T cells after erlotinib treatment, we used an *in vivo* labeling approach to distinguish circulating and parenchymal lung T cells from tumor-bearing mice left untreated or treated with erlotinib for 2 weeks [31]. CD4^+^ and CD8^+^ T cells in the lungs were further classified as naïve or effector based on their expression of molecules involved in lymphocyte migration (e.g. CD62L) necessary for T cell entry into lymph nodes through high endothelial venules [32] and molecules involved in lymphocyte adhesion (e.g. CD44) required to enter sites of inflamed peripheral tissues [33], where interaction with target antigens can occur. Naïve CD4^+^ and CD8^+^ T cells, defined as CD62L^high^ CD44^low^ were unchanged after erlotinib treatment (**Figure 2C**). Conversely, percentages of CD62L^low^ CD44^high^ effector CD4^+^ and CD8^+^ T cells were significantly increased after treatment (**Figure 2D**), suggesting that erlotinib treatment leads to increased effector T cells in the tumor microenvironment. In order to further characterize lung CD4^+^ and CD8^+^ T lymphocytes, we isolated lung-resident CD4^+^ and CD8^+^ T cells and performed RNA sequencing to query their gene expression profiles. As predicted, we detected abundant expression of T cell lineage markers *Cd3e, Cd4, Cd8a* and *Cd8b* in the relevant cell populations that was unchanged by erlotinib treatment (**Supplementary Figures 2B&C**). In addition, we found that T cells from untreated tumors and erlotinib treated tumors have similar levels of expression of the T-cell co-stimulatory molecules *Cd28, Cd27* and *Icos* (**Supplementary Figures 2B&C**). Ingenuity Pathway Analysis (IPA) revealed leukocyte extravasation signaling and agranulocyte adhesion and diapedesis (extravasation) amongst the top ten pathways that changed significantly after erlotinib treatment suggesting that erlotinib treatment modulates lymphocyte properties related to movement and migration (**Supplementary Table 1**).

Next, using cytokine arrays, we measured the protein level of various cytokines and chemokines from whole lung lysates of untreated and treated tumors. Cytokines were categorized based on their ability to induce T cell recruitment, differentiation or function. We found that the T cell chemoattractant CCL5 increased after erlotinib treatment while others such as CXCL10, CXCL9 and CXCL11 did not change (**Figure 2E)**. Cytokines promoting T_H2_ or regulatory T cell responses including IL-4, IL-5, IL-10 and IL-27 were lower after treatment (**Figure 2F**) while the levels of pro-inflammatory cytokines (IFN-γ, IL-12p70, IL-17 and IL-23) that may mediate T_H1_ T cell responses were largely unchanged. Overall, these data suggest that erlotinib promotes the recruitment of inflammatory T cells into the lung tumor microenvironment.

### Tumor regression mediated by erlotinib indirectly leads to the changes in the immune microenvironment

We further questioned whether the effect of erlotinib on the tumor microenvironment was a direct consequence of the TKI or an indirect result of drug-induced tumor regression. To address this question, we leveraged the inducible nature of our model system and removed doxycycline from the diet of tumor-bearing EGFR^L858R^ mice for 2 weeks. Doxycycline withdrawal turns off the transgene initiating rapid tumor cell death similar to that observed with erlotinib [30], (**Figure 3A and Supplementary Figures 3A and 3B**). As is the case with erlotinib, we saw an increase in the percentage of CD4^+^ and CD8^+^ T cells in the lungs of these models (**Figure 3B, Supplementary Figures 3C and D**). Dox withdrawal had a more profound effect on T_regs_ which decreased significantly following oncogene de-induction (along with a corresponding decrease in the T_reg_/CD8 ratio) compared to what was observed with erlotinib treatment (**Figures 3C and 3D**). To further explore whether tumor regression, and not erlotinib directly, was causing the observed changes in the immune microenvironment, we studied mice with *EGFR* mutant lung cancer induced by expression of the EGFR^L858R+T790M^ mutant that is unresponsive to erlotinib treatment (**Supplementary Figures 3A and 3B**) [34]. Following erlotinib treatment of L+T tumor-bearing mice we did not observe changes in the immune microenvironment (**Figures 3B, 3C&3D**). We also treated mono-transgenic healthy littermates with erlotinib for two weeks as an alternative approach to query whether the inhibitor exerts non-specific effects on immune cells and observed no differences in the immune microenvironment between erlotinib treated or untreated lungs (**Supplementary Figures 3E and 3F**). These results lead us to conclude that the changes in the immune microenvironment are not a result of a direct effect of erlotinib on immune cells but rather a consequence of the process of tumor regression itself.

**Figure 3.**
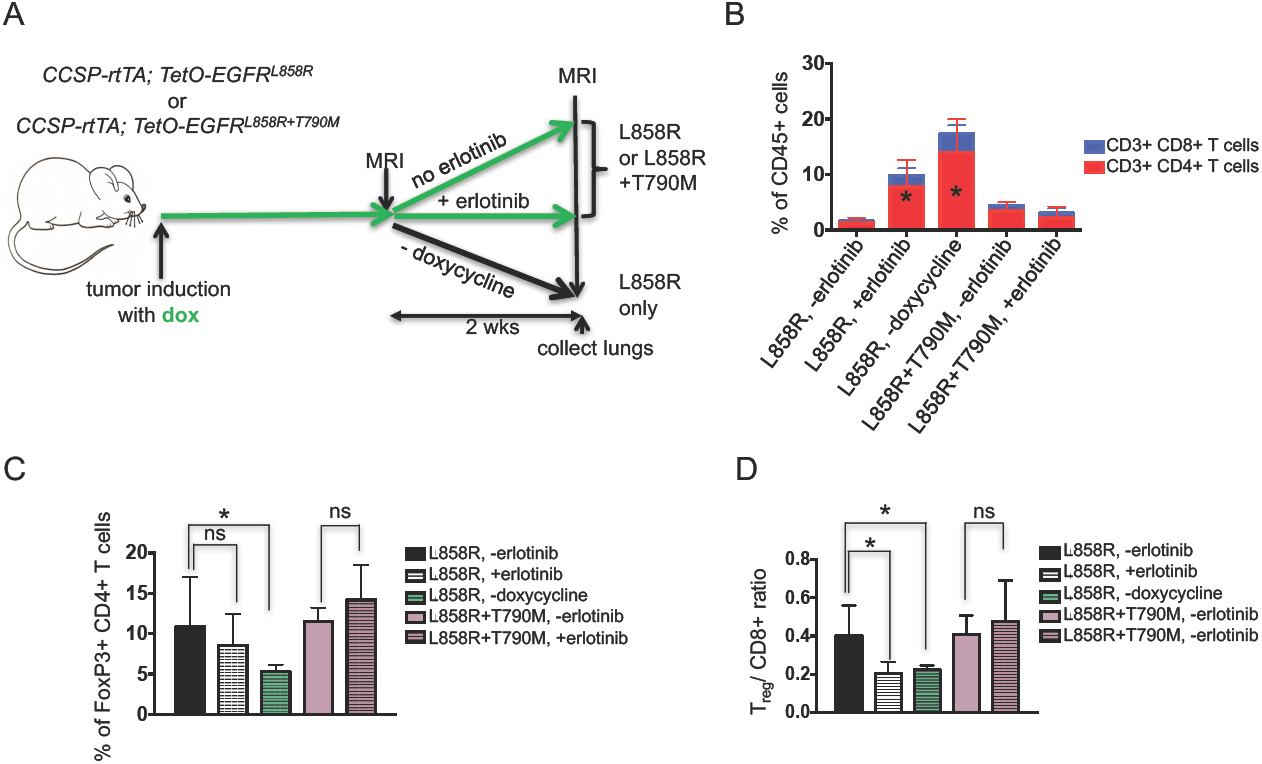
Changes in T cells in the immune microenvironment are due to tumor regression. (**A**) Experimental outline of tumor induction and erlotinib treatment. *CCSP-rtTA; TetOEGFR^L858R^* or *CCSP-rtTA; TetO-EGFR^L858R+T790M^* mice and littermate controls on a doxycycline diet (green arrow) were treated with erlotinib or left untreated for 2 weeks or taken off doxycycline diet. Infiltrating immune cells were analyzed by flow cytometry. Quantification of (**B**) CD4 and CD8 T cells, (**C**) FoxP3 positive CD4 T cells and (**D**) the T_reg_/ CD8 ratio in lungs of tumor bearing *CCSP-rtTA; TetO-EGFR^L858R^* or *CCSP-rtTA; TetO-EGFR^L858R+T790M^* mice in the absence (-) and presence (+) of erlotinib for 2 weeks or after doxycycline withdrawal. Data are from three independent experiments, (n=4-6 mice per group) *, P < 0.05 in student t test.

To further study whether erlotinib directly affects tumor-infiltrating T cells we used *in vivo* labeling to distinguish circulating (i.e., cells in the vasculature) and parenchymal lung T cells followed by flow cytometry analysis. Notably, erlotinib treatment led to an increase in the absolute number of T cells present in the lung epithelium compared to untreated tumor-bearing lungs (**Figure 4A**). This translated into a 4-fold increase in CD4^+^ T cells and 2-fold increase in CD8^+^ T cells (**Figure 4B).** This difference was not as prominent in the circulating T cells collected from the mouse lungs (**Supplementary Figures 4A&4B**). Interestingly, the lung CD4^+^ and CD8^+^ T–cells showed decreased Ki-67 positivity upon erlotinib treatment suggesting that the increased number of these cells was not due to increased proliferation following erlotinib treatment (**Figure 4C**). Similarly, proliferation of circulating CD4^+^ but not CD8^+^ T cells in the lungs was also decreased upon treatment (data not shown). Co-immunofluorescent staining of lung sections with antibodies against CD3 and Ki-67 showed a similar trend (**Figures 4D and E**). Analogous findings were observed in samples from mice following doxycycline withdrawal supporting the possibility that the decrease in T cell proliferation is an indirect effect of the tumor regression rather than a direct effect of erlotinib on the T-cells (**Supplementary Figure 4C**).

**Figure 4.**
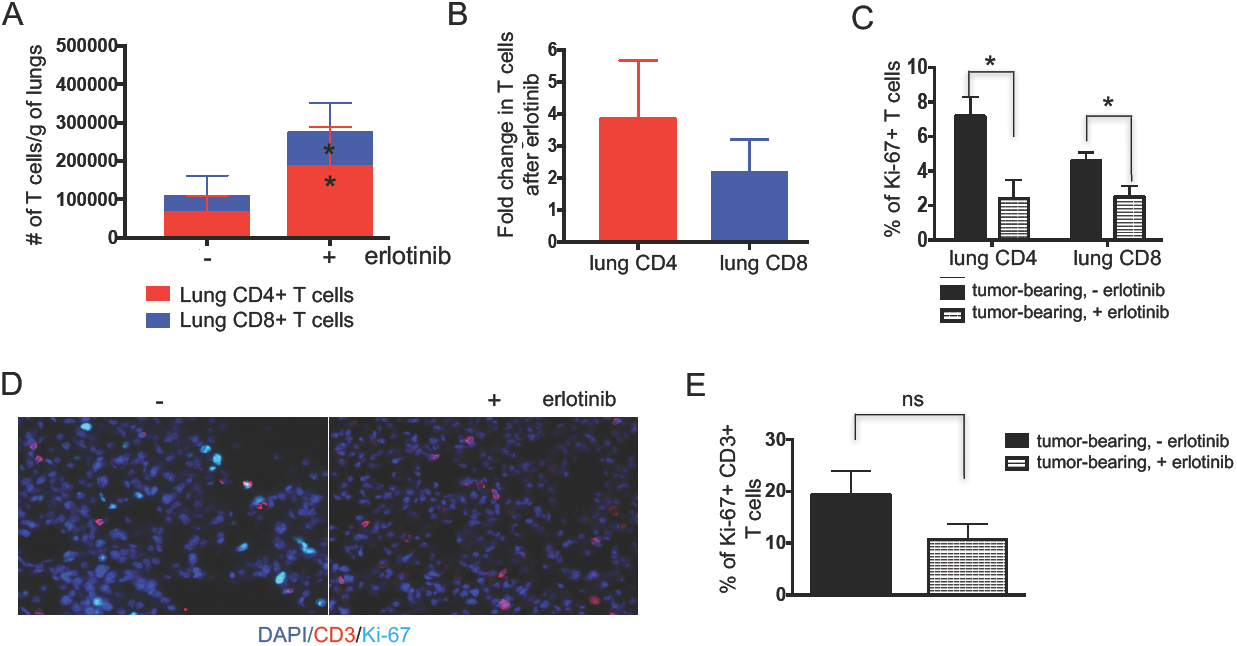
Erlotinib-mediated tumor regression increases lung T cells. (**A**) Absolute number and (**B**) Fold change in number of parenchyma lung CD4 and CD8 T cells of tumor bearing *CCSP-rtTA; TetO-EGFR^L858R^* mice in the absence (-) and presence (+) of erlotinib for 2 weeks. Quantification of (**C**) Ki-67+ CD4 and CD8 T cells of tumor bearing *CCSP-rtTA; TetOEGFR^L858R^* mice in the absence (-) and presence (+) of erlotinib for 2 weeks. (**D**) Immunofluorescent (IF) stain and (**E**) quantification of CD3 T cells (red) and Ki-67 positive cells (Cyan) in lungs of tumor bearing *CCSP-rtTA; TetO-EGFR^L858R^* mice in the absence (-) and presence (+) of erlotinib for 2 weeks. Nuclei were counterstained with Dapi (blue). Data are obtained from three independent experiments, (n=4-6 mice per group) *, P < 0.05 in student t test.

To further confirm that erlotinib did not act directly on T cells, we evaluated its effect on T-cell proliferation by performing CFSE staining (**Supplementary Figures 5A and 5B**) of 10 μM erlotinib and DMSO-treated T cells isolated from the spleen and lungs of tumor-bearing mice. We found that erlotinib, even at this high concentration, did not alter T cell proliferation *in vitro* (**Figures 5A, B and Supplementary Figure 5C**). We also tested the effects of this TKI on T cells after LCMV infection *in vivo* (**Figure 5C**) and found no effect on the abundance of CD44^+^ activated CD4^+^ or CD8^+^ T cells with erlotinib treatment (**Figures 5D&E**). In addition, we did not observe a significant difference in Ki67^+^ CD4^+^ or CD8^+^ T cells between erlotinib and vehicle treated mice suggesting that erlotinib does not affect proliferation of these cells directly (**Figure 5F**).

**Figure 5.**
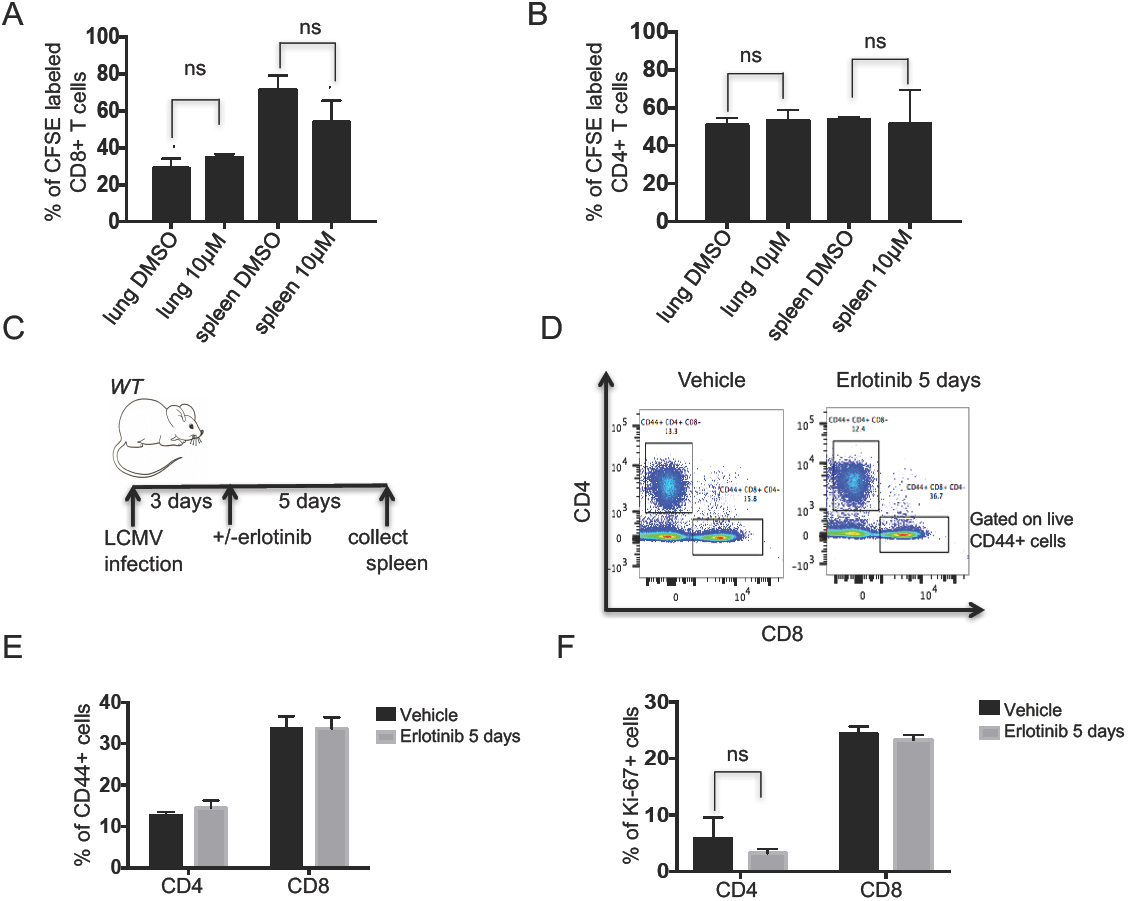
Erlotinib does not diminish T cell proliferation *in vitro* or *in vivo*. Quantification of erlotinib-treated (**A**) CD8 and (**B**) CD4 T cells isolated using magnetic beads from lungs and spleens of tumor bearing *CCSP-rtTA; TetO-EGFR^L858R+T790M^* and labeled with CFSE. The proportion of dividing cells was assessed 120 hours after 10μm erlotinib or DMSO treatment based on CFSE dilution. (**C**) Experimental layout of control, non-tumor bearing *CCSP-rtTA; TetO-EGFR^L858R^* mice infected with LCMV for 8 days with intervening daily administration of erlotinib or vehicle for 5 days. Splenic T cells were collected and analyzed by flow cytometry. (**D**) Representative FACS plot showing the percentage of CD44+ CD4+ or CD44+ CD8+ T cells and quantification of (**E**) CD44+ CD4+ or CD44+ CD8+ T cells. (**F**) Ki-67+ CD4+ or Ki-67+ CD8+ T cells from vehicle or erlotinib treated LCMV infected mice. *, P < 0.05 in student t test.

### Erlotinib treatment leads to increased maturation of myeloid cells

Our cytokine array analysis showed an overall decrease in both myeloid cell-inducing cytokines (**Supplementary Figure 6A**) as well as some chemokines associated with recruitment of myeloid cells (**Supplementary Figure 6B)** after erlotinib treatment. Given these findings, we decided to determine how TKI treatment affected tumor-associated myeloid cells whose presence and functionality may be affected by these cytokines or chemokines. First, we investigated the proportions of myeloid cell populations following erlotinib treatment. Specifically, we measured the percentage of alveolar and interstitial macrophages, neutrophils and dendritic cells (**Figure 6A**). As observed by others [35], there was a prominent expansion of alveolar macrophages (AM) in tumor-bearing mouse lungs and this cell population was significantly decreased after erlotinib treatment (**Figure 6A**) likely due to decreased proliferation of those cells as shown by a lower percentage of Ki-67^+^ positivity in that population after TKI treatment (**Supplementary Figure 6C**). In direct opposition to the pattern observed with AMs, interstitial macrophages and neutrophils were decreased in tumor-bearing lungs compared to controls and increased after erlotinib treatment (**Figure 6A**).

**Figure 6.**
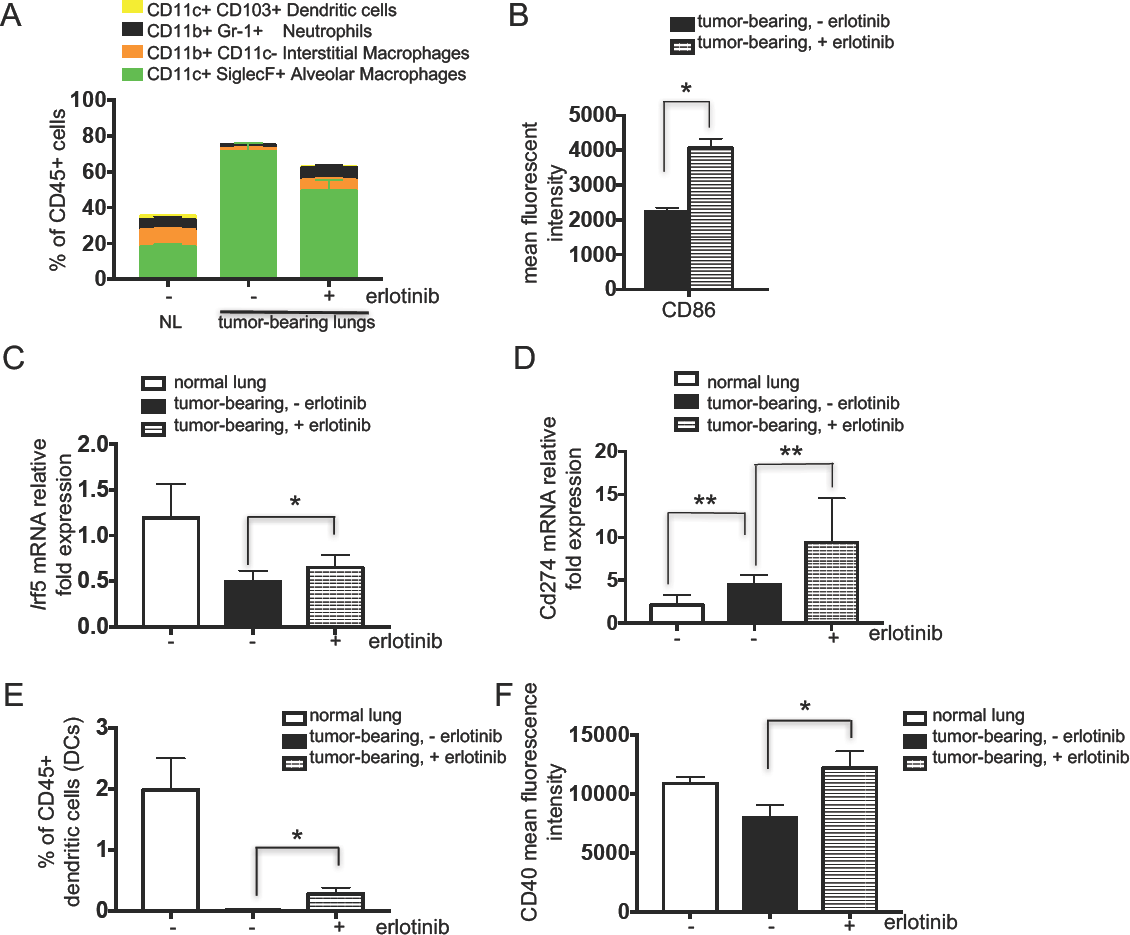
Erlotinib decreases alveolar macrophages and mediates a macrophage phenotypic switch indicative of an improved maturation. Quantification of (**A**) myeloid cell populations, (**B**) mean fluorescent intensity of the co-stimulatory molecule, CD86 in alveolar macrophages (AMs), (**C**) Irf5 and (**D**) Pd-l1 mRNA expression in AMs (**E**) Dendritic cells (DCs) and (**F**) CD40 MFI on DCs from lungs of control (normal) and tumor bearing *CCSP-rtTA; TetOEGFR^L858R^* mice in the absence (-) and presence (+) of erlotinib for 2 weeks. Data are obtained from three independent experiments, (n=4-6 mice per group) *, P < 0.05 in unpaired student t test.

Pulmonary AMs serve diverse roles in defense against pathogens in the respiratory tract. In addition to their well-established phagocytic roles and microbicidal functions [36], they also initiate pro-inflammatory responses through secretion of cytokines, which can stimulate T helper type 1 (T_H1_) T cell responses or anti-inflammatory responses through secretion of IL-10 [37]. Finally, AMs have been described as poor antigen presenting cells, due to low expression of the co-stimulatory molecules CD80 and CD86 [38]. We observed an increase in the mean fluorescence intensity of CD86 on AMs suggesting a mature antigen presenting phenotype (**Figure 6B**). Further supporting a switch in the macrophages to a pro-inflammatory phenotype, *Irf5* expression was increased in AMs isolated from erlotinib-treated lungs (**Figure 6C**). High expression of *Irf5* has been shown to be characteristic of pro-inflammatory M1 macrophages, which are potent promoters of T_H1_ T cell responses [39]. The levels of expression of M2 macrophage markers such as Chitinase-like 3 or MRC-1 were unchanged in lung tumors compared to healthy lungs or after erlotinib treatment (data not shown). Interestingly, gene expression of *Cxcl2* increased in AMs after erlotinib treatment (**Supplementary Figure 6D**). This could potentially explain the increased neutrophils observed in TKI-treated lungs (**Figure 6A**). These results suggest that erlotinib treatment triggers an inflammatory response in AMs.

Since a decrease in CD8^+^ T cell responses can be mediated by immune checkpoint ligands, like PD-Ligand 1 (PD-L1 or B7H1), we investigated whether the levels of this molecule were modulated by erlotinib. We found increased *Cd274* (the gene encoding PD-L1) expression on AMs after erlotinib treatment (**Figure 6D**), perhaps as a consequence of an adaptive immune response triggered by the inflammatory microenvironment induced by erlotinib. Moreover, IFN-γ secreted by activated effector T cells, described earlier, has been shown to induce PD-L1 in mouse models [40]. However, we did not observe a significant difference in expression of *Cd274* on Epcam+ cells from normal lungs compared to cells from tumor bearing or erlotinib treated lungs (**Supplementary Figure 6E**). Dendritic cells were notably absent in tumor-bearing untreated lungs compared to their healthy lungs counterparts. Surprisingly, there was a significant increase in CD103+ dendritic cells after erlotinib treatment (**Figure 6E**). These dendritic cells exhibited higher expression of CD40 (**Figure 6F**), a co-stimulatory molecule required for activation of antigen presenting cells. Taken together, these results suggest that the antigen-presenting cells present in the tumor microenvironment after erlotinib treatment exhibit a more mature phenotype than cells in untreated tumors that may render them capable of driving tumor antigen specific responses.

## Discussion

In this study, we investigated the changes that occur within the immune microenvironment in a mouse model of *EGFR* mutant lung cancer after treatment with the TKI erlotinib. We found that erlotinib treatment led to the establishment of an inflammatory immune microenvironment in the lungs. Importantly, these changes were not due to a direct effect of the TKIs on cells in the immune microenvironment but rather they were stimulated by the process of tumor regression itself.

Given the increasing interest in combining targeted therapies and immunotherapies, efforts to study the consequences of targeted therapies on the tumor immune microenvironment are growing [43]. Our findings demonstrating that erlotinib is partially immunostimulatory are consistent with observations made with other targeted therapies. Studies of the BRAF inhibitor vemurafenib in a mouse model of *Braf* mutant and *Pten* deficient melanoma showed increased cytokine producing T cells in tumors following kinase inhibitor treatment [41, 44]. Similarly, activated CD8 cells were also more abundant in a *Kit* mutant gastrointestinal stromal tumor (GIST) model after imatinib treatment [45]. These indications of immune activation were counterbalanced by increased levels of the immune inhibitory ligand PD-L1 observed in both the melanoma and lung models (**Figure 6E**) following kinase inhibitor treatment indicating that elements of immunesuppression are still present after treatment with these targeted agents. These findings highlight some of the similarities of the effects of kinase inhibitors on tumors in mouse models, however there are also some unique consequences of different inhibitors. For example, in the GIST model imatinib also leads to depletion of T_regs_ in addition to leading to an increase in CD8 effector T cells [45].

There are several ways in which targeted therapies may be affecting immune cells. They could either be acting directly via on-target or off-target activities on immune cells present in the tumor. Alternatively, the changes could be an indirect consequence of the biological effects (e.g induction of apoptosis) of targeted therapies. Indeed, forms of cell death, like necrosis, have long been recognized as having potentially immunogenic consequences, and data suggest that apoptosis could also have immunological effects [46]. In support of this, our study provides evidence that the TKI erlotinib itself does not act directly on immune cells in the tumor microenvironment but rather changes in immune infiltrates result indirectly from the process of tumor regression. First, we found that in a mouse model of erlotinib-resistant lung cancer in which tumors do not regress upon treatment with the TKI, low numbers and functionally impaired CD4 and CD8 lymphocytes are found similar to untreated tumors even following TKI treatment. Second, erlotinib did not affect the proportion of lymphocytes in the lungs of healthy non-tumor bearing mice. Third, erlotinib treatment of lymphocytes isolated from tumor-bearing mouse lungs or from spleen does not affect their proliferation or activation. Others have shown that erlotinib does inhibit the proliferation of T cells isolated from mouse lymph nodes [27]. It is possible that these differences are due to the different biological contexts examined, namely lung or spleen cells from tumor-bearing or LCMV-infected mice as opposed to T cells from wild-type lymph nodes.

Despite changes towards an immunostimulatory microenvironment after erlotinib treatment several findings indicated that the tumors retained some immunosuppressive properties including abundant regulatory T cells (**Figure 1C**) and increased levels of PD-L1 (**Figure 6D**). While the T_regs_ may be indicative of immunosuppression persistent after erlotinib, the cells may also be playing a role in tissue repair after inflammation [47]. Whether targeting these elements of immunosuppression would be an effective strategy to slow tumor growth is currently unknown and actively being investigated. Such studies could include direct targeting of T_regs_ either by using clinically approved antibodies such as ipilimumab (anti-CTLA-4) that can deplete T_regs_ [48] or, in genetically engineered mouse models, by ablating T_regs_ [49]. PD-1 axis inhibitors have been shown to modestly prolong survival of mice with EGFR mutant lung cancer [20], however, whether in combination with erlotinib this translates into improved survival and/or delays the emergence of resistance is unknown. In patients, the response rate of EGFR mutant tumors to PD-1 or PD-L1 blockade is ∼10% and therefore lower than in NSCLC as a whole (RR ∼20%) potentially due to the lower immunogenicity of the tumors mainly arising in former/never smokers and having a low mutational background [14, 16, 19, 50]. Large studies of TKIs in combination with checkpoint inhibitors have not been conducted to date in part due to concerns regarding the toxicity of these combinations. However, in a small study of erlotinib in combination with nivolumab, the combination was well-tolerated and the response rate to the combination in the refractory setting was 15% suggesting that some patients benefit from these agents [21].

The low response rate to PD-1 inhibitors in human EGFR mutant lung cancers and the relatively modest effects of these drugs alone in EGFR mutant tumors in mice suggest that leveraging the immune system in these tumors is challenging. It has been established that lung tumors in genetically engineered mouse models, including the EGFR^L858R^ model we used, have a significantly lower frequency of nonsynonymous mutations compared to human lung adenocarcinomas [51, 52]. The low frequency of somatic mutations that arise during tumor development in these models lead to the generation of few neoantigens to induce T cell responses. This may explain the lack of a strong T cell-mediated immune response in this tumor model [53]. Future studies aimed to study antigen-specific T cell responses in new systems that express model antigens and/or have higher mutation burdens more reflective of human lung cancer are ongoing. An alternative but not mutually exclusive possibility, is that multiple immunosuppressive pathways active in the tumors need to be simultaneously inhibited to engage the immune system. This is supported by our data showing that suppressive cytokines increase in abundance and that T_regs_ are unchanged in EGFR^L858R^-induced tumors following erlotinib treatment. The extent to which these signals play a role in tumorigenesis and need to be reversed for tumor regression is still poorly understood.

Altogether, our findings lay the foundation for understanding the tumor immune microenvironment in the tumor regression process mediated by TKIs. The absence of a complete reversal of the immunosuppressive environment present in lung tumors after treatment with erlotinib provides an opportunity for exploring treatment combinations that target both the immune microenvironment and tumor cells.

## Methods

### Transgenic Mice

*CCSP-rtTA; TetO-EGFR^L858R^* mice were previously described [30]. Mice were fed chow containing doxycycline (625 ppm) obtained from Harlan-Tekland. The animals were housed in a pathogen-free facility according to approved guidelines from the Yale University Institutional Animal Care and Use Committee (IACUC).

### In vivo treatment with Erlotinib

Erlotinib was purchased and purified at the organic synthesis core facility at Memorial Sloan Kettering Cancer Center (MSKCC), dissolved in 0.5% methylcellulose and administered intraperitoneally at 25mg/kg, 5 days a week. Mice were euthanized by CO_2_ asphyxiation.

### Magnetic resonance imaging

Magnetic resonance images of isofluroane-anesthetized mice were collected using a mini-4T horizontal-bore spectrometer (Bruker AVANCE). Throughout data collection, each animal was anesthetized on a steady flow of isofluroane and oxygen (2-2.5% v/v) and core-body temperature was maintained at 37±1°C. Imaging parameters were optimized to effectively discriminate between healthy lung and areas with tumor. Tumor burden in each animal was quantified by calculating the volume of visible lung opacities in every image sequence using the BioImage Suite software [54].

### Tumor digestion

Lungs from normal, untreated, tumor-bearing or treated mice were mechanically digested and incubated in HBSS with 0.5mg/ml collagenase IV and 1ug/ml DNase 1 at 37 degrees for 1 hour after which the solution was filtered using a 70μm cell strainer. The resulting single cell suspension was incubated in ACK lysis buffer for 5 minutes to lyse red blood cells.

### Flow cytometry and cell sorting

Single cell suspensions of lung tumors or splenocytes were resuspended in FACS buffer (PBS+1%FBS). Cells were then incubated with anti-Fc receptor antibody (clone 2.4G2) on ice for 15 min s followed immediately by staining with respective surface antibodies for 30 minutes. For intracellular cytokines, T cells were stimulated with PMA/ionomycin (Sigma Aldrich) and Brefeldin A for 5 hours at 37 degrees. The cells were stained first with surface antibodies then fixed in Cytofix/Cytokperm buffer (BD Biosciences) followed by staining with antibodies to detect proteins present in intracellular compartments. FoxP3 staining was done in a similar manner. Samples were acquired on LSRII flow cytometer and analyzed with Flowjo. Cells were sorted on the BD FACS Aria at the Yale Cell Sorter Core facility. Cells were sorted based on the expression of the following markers: CD4 T cells: CD45+/CD3+/CD4+, CD8 T cells: CD45+/CD3+/CD8+, Alveolar macrophages: CD45+/ CD11c+/SiglecF+, Tumor epithelial cells: CD45-/CD11c-Epcam+.

### In vivo labeling of immune cells

Mice were retro-orbitally injected with 3μg of biotin-conjugated CD45 (clone 30-F11) for 5 minutes, immediately after which animals were sacrificed. Lung tissue was collected, processed and stained as described above.

### T cell proliferation assay

Splenocytes and single cells suspension were collected from spleen or lungs of tumor bearing mice. T cells were enriched using a purified antibody cocktail consisting IA/I-E, B220 and F4/80. Purified cells were loaded with 5μM CFSE at room temperature for 15 minutes in the dark. T cells mixed with anti-CD28 were seeded on CD3 coated plates followed by treatment with 10μM Erlotinib or DMSO for 5 days. Proliferation was determined by CFSE dilution using flow cytometry.

### Histology, Immunofluorescent Assay and Cell Quantification

Lung tissue from normal, tumor bearing untreated and treated animals was collected after sacrifice, fixed overnight in 4% paraformaldehyde and rehydrated in 70% ethanol until submission for paraffin embedding and sectioning at the Yale Pathology Tissue Services. Sections were stained with hematoxylin and eosin, CD3 (Spring Biosciences; 1:150), EGFR^L858R^ (Cell Signaling; 1:400), FoxP3 APC-conjugated (eBioscience; 1:50), Ki-67 (BioLegend; 1:50) and Cytokeratin 7 (Abcam; 1:300) antibodies. Positive cells in a 40X field of view were manually counted using a plugin for ImageJ called Cell Counter. At least three representative tissue locations were used to quantify and values were averaged for each mouse.

### RNA extraction, purification and quantitative real-time RT-PCR

For RNA extraction and purification, the Arcturus PicoPure RNA isolation kit was used according to manufacturer’s instruction and cDNA was synthesized using the SuperScript II Reverse Transcriptase from Invitrogen. Quantitative real-time PCR was performed using the Taqman assay (Invitrogen). C_t_ values were recorded and relative gene expression was determined using the ΔΔC_t_ method.

### RNA Sequencing and Gene expression data

RNA sequencing was performed using the illumina HiSeq 2000 platform through the Yale Stem Cell Center Genomics Core facility. R1 reads from each paired-end reads were aligned to mouse genome (version mm10) using bowtie2 [55] in local mode, followed by annotation of counts to each gene by gencode (version M10) [56]. Differential expression in each cell type between experimental conditions was performed with the DESeq2 [57] R package.

### Ingenuity Pathway Analysis

Enrichment analyses of canonical pathways were performed with Ingenuity Pathway Analysis (IPA, Ingenuity Systems). Genes with an adjusted P value lower than 0.05 were included and Ingenuity Knowledge Base (Genes Only) was used as reference set for the analyses.

## Acknowledgements

Sequencing service was conducted at Yale Stem Cell Center Genomics Core facility, which is supported by the Connecticut Regenerative Medicine Research Fund and the Li Ka Shing Foundation. We thank Dr. Nikhil Joshi for critical reading of the manuscript.

## Grant Support

This work was supported by NIH/NCI grants R01CA195720 (KP and SK) and P50CA196530 (KP), a Yale Cancer Center Co-Pilot award (KP and SG), the National Science Foundation Graduate Research Fellowship Program (GRFP) (D. Ayeni) and the National Cancer Institute (NCI) Ruth L. Kirschstein National Research Service Award (NRSA) Pre-doctoral fellowship (D. Ayeni).

## Supplementary Figure Legends

**Supplementary Figure 1.**
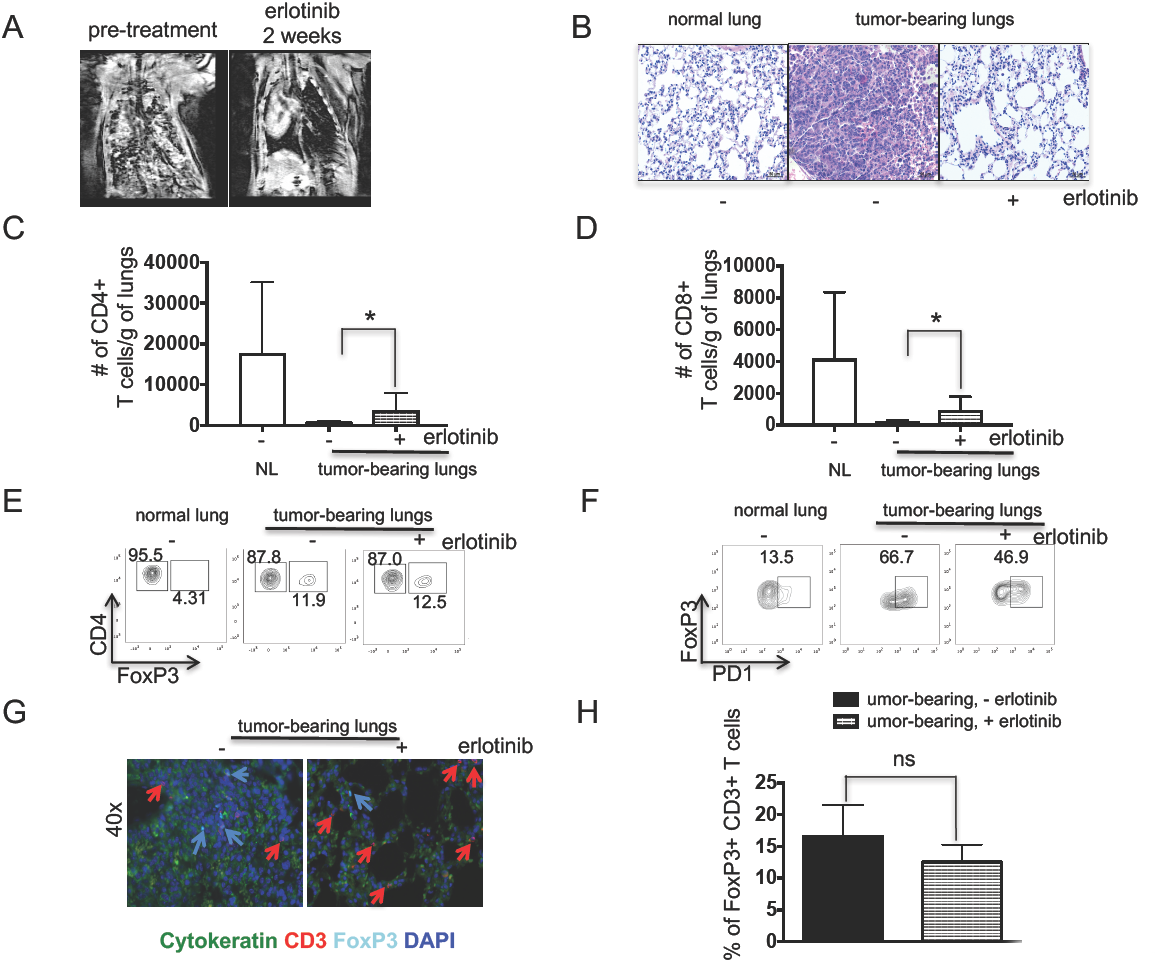
MRI images, histology and representative flow cytometry plots of normal or tumor-bearing lungs before and after erlotinib. (**A**) Coronal images of *CCSP-rtTA; TetO-EGFR*^*L858R*^ mouse lungs before (left panel) and after (right panel) treatment with erlotinib. (**B**) Hematoxylin and eosin (H&E) stain of lungs from control (normal) and tumor bearing *CCSP-rtTA; TetO-EGFR*^L858R^ mice in the absence (−) and presence (+) of erlotinib for 2 weeks. Bar: 50μm. Absolute number of (**C**) CD4 and (**D**) CD8 T cells normalized to weight of lungs of control (normal) and tumor bearing *CCSP-rtTA; TetO-EGFR*^*L858R*^ mice in the absence (−) and presence (+) of erlotinib for 2 weeks. Representative FACS plot showing percentage of (**E**) FoxP3+ and FoxP3- CD4+ T cells (**F**) PD1+ FoxP3+ T cells. (**G**) Immunofluorescence (IF) stain of lung epithelial cells (green), CD3 T cells (red) and FoxP3 Tregs (Cyan). Nuclei were counterstained with Dapi (blue) (**H**) Quantification of FoxP3+ CD3 T cells in lung tumor bearing *CCSP-rtTA; TetO-EGFR*^*L858R*^ mice in the absence (−) and presence (+) of erlotinib for 2 weeks stained by IF.

**Supplementary Figure 2.**
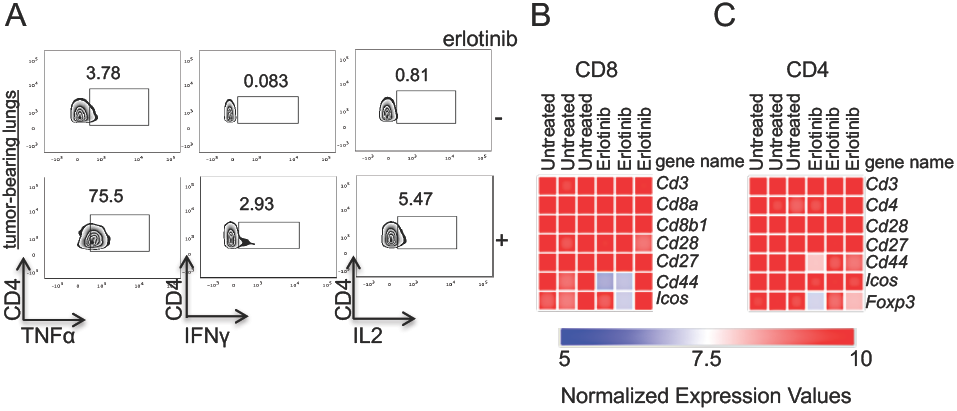
Representative flow cytometry plots of cytokine producing T cells and gene expression profile of T cells isolated from tumor-bearing lungs before and after erlotinib. Representative FACS plots showing the percentage of (**A**) TNF-α+, IFN-γ+, and IL-2+ CD4 T cells. (**B**) CD8 and (**C**) CD4 T cells isolated from *CCSP-rtTA; TetO-EGFR*^*L858R*^ tumor bearing mice untreated or treated with erlotinib. Heatmap was generated using normalized expression values.

**Supplementary Figure 3.**
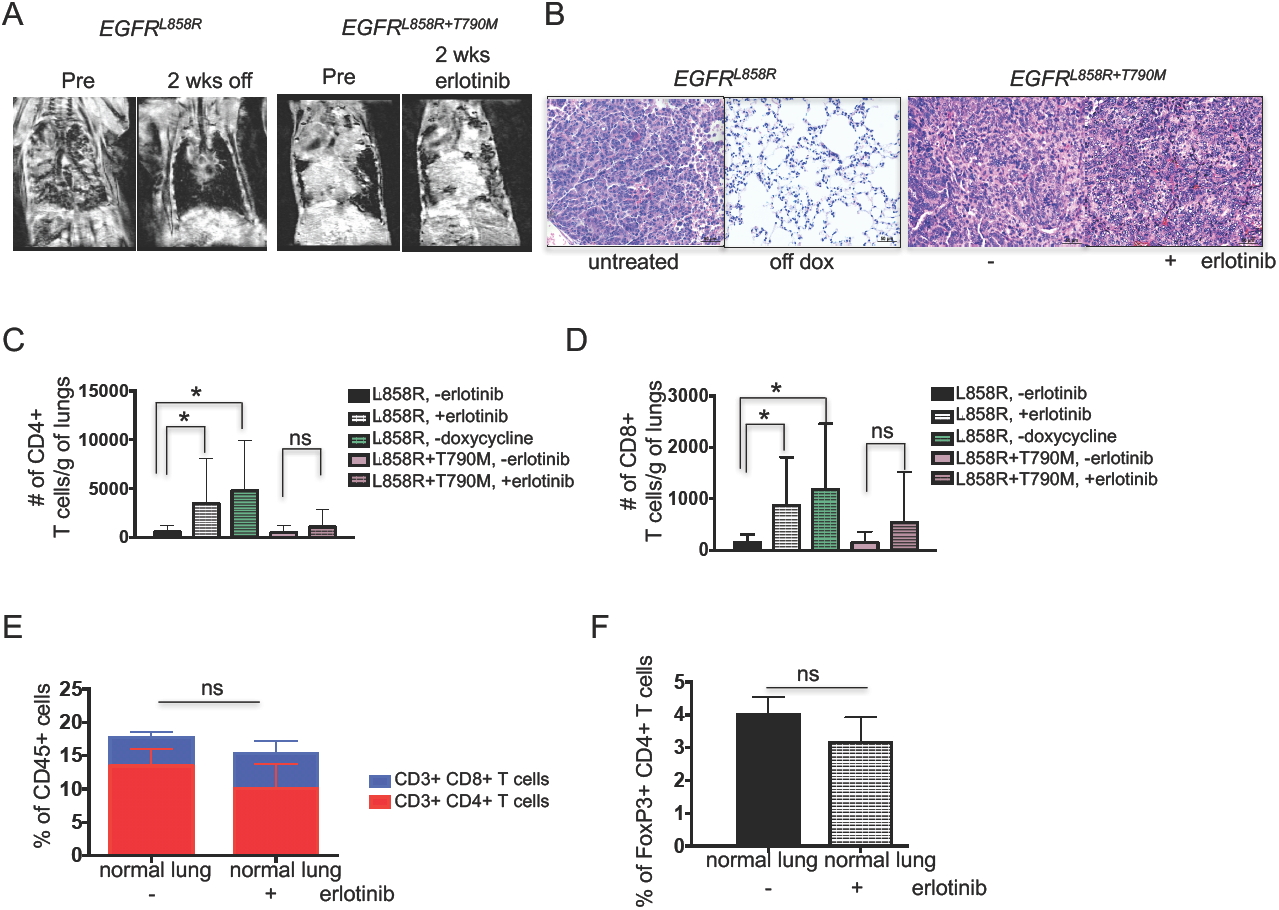
MRI images, histology and representative flow cytometry plots of erlotinib sensitive and resistant tumors. (**A**) Coronal images of: *CCSP-rtTA; TetO-EGFR*^*L858R*^ mouse lungs before (left panel) and after (right panel) cessation of doxycyline and *CCSP-rtTA; TetO-EGFR*^*L858R+T790M*^ mouse lungs before (left panel) and after (right panel) treatment with erlotinib. (**B**) Hematoxylin and eosin (H&E) stain of lungs from tumor bearing: *CCSP-rtTA; TetO-EGFR*^*L858R*^ untreated or taken off doxycycline diet for 2 weeks and *CCSP-rtTA; TetO-EGFR*^*L858R+T70M*^ mice in the absence (−) and presence (+) of erlotinib for 2 weeks. Bar: 50μm. Absolute number of (**C**) CD4 and (**D**) CD8 T cells normalized to weight of lungs of tumor bearing *CCSP-rtTA; TetO-EGFR*^*L858R*^ or *CCSP-rtTA; TetO-EGFR*^*L858R+T790M*^ mice in the absence (−) and presence (+) of erlotinib for 2 weeks or taken off doxycycline diet. Data are obtained from three independent experiments, (n=4−6 mice per group) *, P < 0.05 in unpaired student t test. Quantification of (**E**) CD4 and CD8 T cells and (**F**) FoxP3 positive CD4 T cells in the lungs of control (normal) mice in the absence (−) and presence (+) of erlotinib for 2 weeks.

**Supplementary Figure 4.**
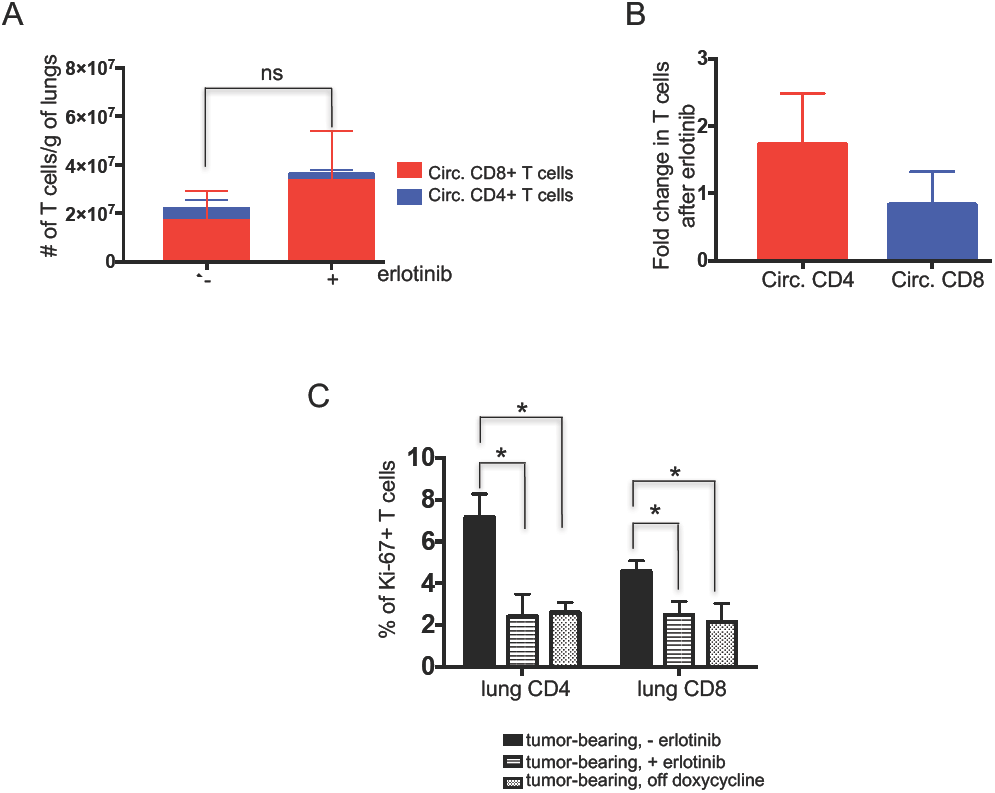
Quantification of circulating and proliferating T cells. (**A**) Absolute number and (**B**) Fold change in number of circulating lung CD4 and CD8 T cells of tumor bearing *CCSP-rtTA; TetO-EGFR*^*L858R*^ mice in the absence (−) and presence (+) of erlotinib for 2 weeks. (**C**) Ki-67+ CD4 and CD8 T cells of tumor bearing *CCSP-rtTA; TetO-EGFR*^*L858R*^ mice in the absence (−) and presence (+) of erlotinib for 2 weeks or mice taken off doxycycline for 2 weeks. *, P < 0.05 in unpaired student t test.

**Supplementary Figure 5.**
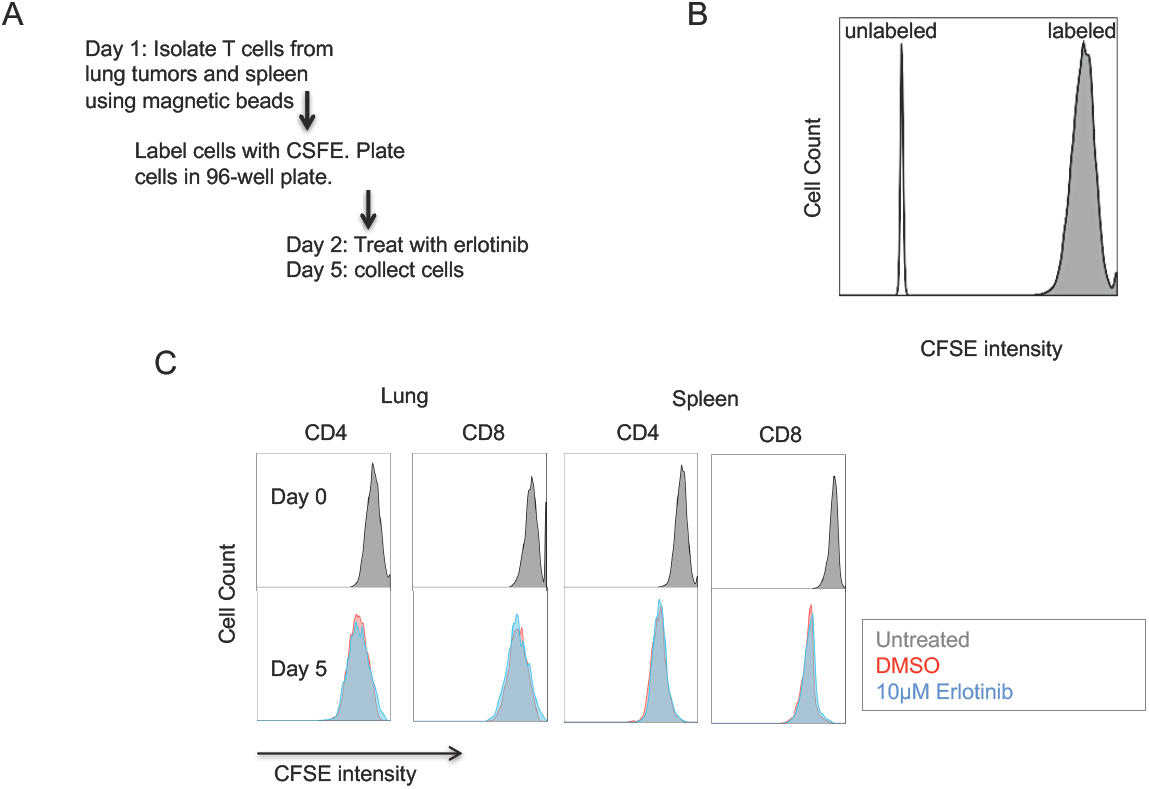
Experimental outline of CFSE labeling and analysis. (**A**) Flow chart for isolation, labeling and treatment of T cells from lungs and spleens of tumor bearing CCSP-rtTA; TetO-EGFRL858R mice. FACS plot showing (**B**) as a control for the technique, unlabeled vs CFSE labeled splenocytes (Day 0) and (**C**) Untreated CD4 and CD8 T cells from lungs and spleens at Day 0 as well as CD4 and CD8 T cells from lungs and spleens treated with 10μm erlotinib or DMSO after 5 days (120 hours).

**Supplementary Figure 6.**
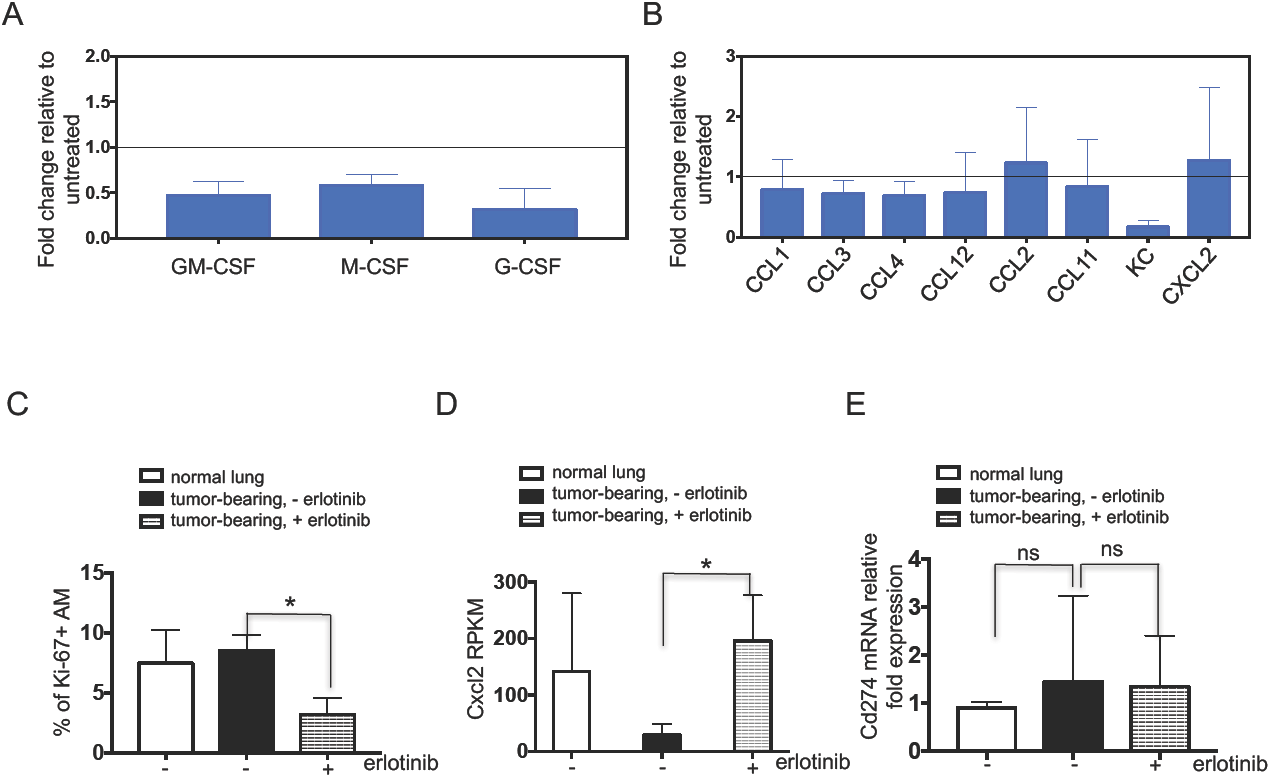
Cytokine profile of myeloid cells in tumor-bearing lungs before and after erlotinib. Graph representing (**A**) cytokine and (**B**) chemokine levels in erlotinib treated whole lung tissue lysate normalized to the levels in untreated tumor bearing lungs from CCSP-rtTA; TetO-EGFRL858R mice. (**C**) Ki-67 positive AMs, (**D**) CXCL2 expression in AMs, (**E**) PD-L1 mRNA expression in Epcam+ tumor cells from lungs of control (normal) and tumor bearing CCSP-rtTA; TetO-EGFRL858R mice in the absence (−) and presence (+) of erlotinib for 2 weeks. *, P < 0.05 in unpaired student t test.

